# Autonomic Nervous System activity correlates with peak experiences induced by DMT and predicts increases in wellbeing

**DOI:** 10.1101/2024.03.19.585567

**Authors:** Valerie Bonnelle, Amanda Feilding, Fernando E. Rosas, David J. Nutt, Robin L. Carhart-Harris, Christopher Timmermann

## Abstract

Non-ordinary states of consciousness induced by psychedelics can be accompanied by so-called ‘peak experiences’, characterized at the emotional level by their intensity and positive valence. These experiences are strong predictors of positive outcomes following psychedelic-assisted therapy, and it is therefore important to better understand their biology. Despite growing evidence that the autonomic nervous system (ANS) plays an important role in mediating emotional experiences, its involvement in the psychedelic experience is poorly understood. The aim of this study was to investigate to what extant changes in the relative influence of the sympathetic (SNS) and parasympathetic nervous systems (PNS) over cardiac activity may reflect the subjective experience induced by the short-acting psychedelic N,N-Dimethyltryptamine (DMT). We derived measures of SNS and PNS activity from the electrocardiogram data of 17 participants (11 males, 6 females, mean age = 33.8 y, SD = 8.3) while they received either DMT or placebo. Results show that the joint influence of SNS and PNS (‘sympatho-vagal coactivation’) over cardiac activity was robustly correlated with participants ratings of ‘Spiritual Experience’ and ‘Insightfulness’ during the DMT experience, while also being related to improved wellbeing scores two weeks after the session. In addition, we found that the state of balance between the two ANS branches (‘sympatho-vagal balance’) before DMT injection predicted scores of ‘Insightfulness’ during the DMT experience. These findings demonstrate the important involvement of the ANS in psychedelic-induced peak experiences and may pave the way to the development of biofeedback-based tools to enhance psychedelic-therapy.

**Significance statement:** Psychedelics can give rise to intense positive subjective experiences - aligned with Maslow’s notion of ‘peak experiences’ - that can have a positive and enduring impact on mental health. Understanding how these experiences relate to peripheral physiology before and during the acute effects of psychedelics is an important object of enquiry, as it may help advance the therapeutic use of these compounds. In this study, we demonstrate that specific peripheral states computed from heart rate activity recordings predicted and correlated with acute peak experiences and increases in wellbeing. These findings have implications for the relationship between peripheral physiology and altered states of consciousness. Moreover, they highlight a putative marker of physiological ‘readiness’ prior the psychedelic experience that could predict therapeutically relevant mechanisms that might be modified to improve mental health outcomes in psychedelic-therapy.

## Introduction

Psychedelics induce significant alterations in perceptual, cognitive, and emotional processing and in scientific studies can act as tools of perturbation for human consciousness (1). Evidence suggests that intense positively valanced self-transcendent experiences (also called ‘peak experiences’) occurring during psychedelic therapy significantly predict positive mental health outcomes (2, 3), such as reductions in depression and anxiety (4), as well as reduced craving in substance abuse disorders (5). Despite the important therapeutic relevance of these experiences, the mechanisms leading to these states remains unclear. In addition, even though great care may be taken to optimize psychedelic therapy to facilitate these experiences (6), attempting to predict the affective features of a psychedelic experience is still challenging.

Contemporary research has largely focused on the effects of psychedelics on the central nervous system, ignoring other biological mechanisms potentially involved in the emotional states induced by these substances. Yet, brain and body are undeniably intrinsically and dynamically linked; perceptions, emotions, and cognitions not only influence, but also respond to the state of the body (7), as intuited by William James over a century ago (8) and emphasized by the embodied approach to cognitive neuroscience (9). The importance played by the Autonomic Nervous System (ANS) – particularly at the cardiac level – in emotional experiences and affective states is becoming increasingly recognized (10–12). Although the centrality and specificity of the autonomic response is still subject to debate, recent work importantly demonstrated that cardiac sympatho-vagal activity may initiate emotional responses preceding neural dynamics (13). To what extent, then, are the intense emotional experiences fostered by psychedelics mediated by their impact on ANS activity and heart function?

The ANS has two main branches: the sympathetic nervous system (SNS) and the parasympathetic nervous system (PNS). The SNS is involved in priming the body and brain for action, triggering a series of physiological changes that are part of the stress response (“fight-or-flight”). The SNS has generally been proposed to be involved in emotional arousal, when it is experienced with positive (e.g., joy, excitement) or negative (e.g., anxiety, hyper-arousal) valence (11, 14). Classic psychedelics (e.g., LSD, psilocybin, mescaline, and DMT) are serotonin-2A (5-HT2A) receptor agonists and exert effects consistent with activation of the SNS, particularly pupillary dilation, increases in heart rate and blood pressure, and increases in plasma stress hormones cortisol and epinephrine (15–18). The PNS on the other hand, is involved in rest and recovery after periods of stress, energy conservation/storage and regulation of bodily functions (19). While parasympathetic activity has often been found to be associated with positive and pro-social emotions (11, 20), the relation between emotional valence and PNS is not straightforward (11). The PNS does not appear to be directly activated by classic psychedelics. Rather, serotonin 2A receptors antagonist ketanserin, known to reduce or even prevent psychoactive effects when administered simultaneously with a classic psychedelic, has been found to cause an increase in parasympathetic activity (18). The PNS might however become active throughout the experience as a homeostatic response to the initial stress response induced by psychedelics, a mechanism known as ‘vagal rebound’ (21, 22). A commonly held view is that the sympathetic and parasympathetic branches of the ANS are subject to reciprocal central control, with increasing activity of one branch associated with decreasing activity of the other. However, it has now been established that autonomic control of dually innervated target organs, such as the heart, cannot adequately be viewed as a continuum extending from parasympathetic to sympathetic dominance (23). Rather, the two autonomic branches can vary reciprocally, independently, or coactively. We refer to this joint influence as ‘sympatho-vagal coactivation’.

Intensely pleasurable experiences are often associated with autonomic manifestations (e.g., ‘chills’), which are accompanied by physiological markers of arousal such as increase heart rate (24). On the other hand, contemplative experiences (e.g., mindfulness meditation), which can also lead to peak experiences, are typically associated with increased parasympathetic activity (25, 26). However, to our knowledge, the interplay between autonomic activity and peak experiences - which share some characteristics of the two types of experiences mentioned above - has never been investigated. Furthermore, if we consider sympathetic tone as reflecting our ability to respond to stress (and/or salience), and parasympathetic tone as our ability to regulate and restore homeostasis after a stress response, it then becomes apparent that a state of balance between the two branches (sympatho-vagal balance), might be beneficial for adaptation, resilience to stress, and emotional flexibility (20, 27). These attributes may play a crucial role in navigating the often challenging emotional states induced by psychedelics (28).

N,N-dimethyltryptamine (DMT), a short-acting serotonergic psychedelic with therapeutic potential (29, 30) offers a unique opportunity to investigate shifts in autonomic activity and their relation to the psychedelic experience as its short duration of action (∼12 minutes) allows drawing more accurate relationships between subjective reports of acute effects, physiological measures, and mental health outcomes. Crucially, it also provides an optimal opportunity to assess how these dynamics relate to peak experiences, and changes in wellbeing. Here, our primary aim was to investigate the complete profiles of SNS and PNS fluctuations throughout the DMT experience, and to evaluate to what extent these profiles may reflect participants’ subjective experience. We hypothesized that 1) starting the experience from a state of greater sympatho-vagal balance may be conducive of peak experiences, and that 2) peak experiences may be associated with sympatho-vagal coactivation (**Figure 1**).

**Figure 1.**
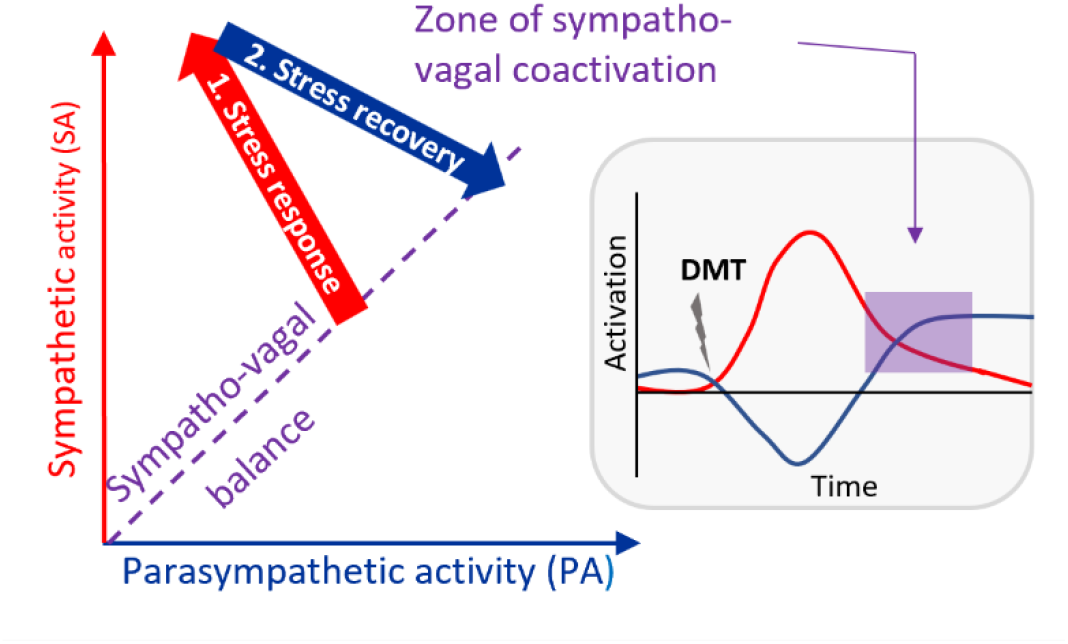
Proposed model for ANS involvement during the psychedelic experience. The hypothesis proposed here is that peak experiences, which are characterised by their intensity and positive balance, are associated with the dual influence of the sympathetic (red) and parasympathetic branches of the ANS (blue) over cardiac activity. Psychedelics may facilitate the induction of this ‘sympatho-vagal coactivation’ by triggering an initial stress response associated with strong sympathetic stimulation and moderate parasympathetic withdrawal, followed by a recovery phase largely mediated by parasympathetic activity increase. We further hypothesised that starting the experience from a state of balance between the SNS and PNS (sympatho-vagal balance) may be optimal in order to return to a balanced, but more activated state, after the initial stress response, which we hypothesize to be optimal for peak experiences.

## Materials and Methods

### Participants and Experimental Procedures

The study was designed as a single-blinded placebo-controlled trial in healthy participants. Experimental sessions consisted of continuous and simultaneous fMRI-EEG resting-state scans which lasted 28 min, with DMT or placebo administered at the end of the 8th min. Participants laid in the scanner with their eyes closed (an eye mask was used to prevent eyes opening), while brain activity and electrocardiograms (ECG) were recorded. Following the scanning procedure, participants were interviewed and completed questionnaires designed to assess the subjective effects experienced during the scan [Visual Analog Scales and validated scales: 11 Dimensions Altered States of Consciousness Questionnaire—ASC-11D (31) and the Mystical Experience Questionnaire—MEQ-30 (32)]. A second session then followed with the same procedure as the initial session, except on this occasion, participants were (audio) cued to verbally rate the subjective intensity of drug effects every minute in real time while in the scanner (**Figure 2**). This article reports the results concerning the ECG data collected during resting-state scans where participants received either DMT or a placebo, and were not interrupted for experience sampling purposes. In total, 20 participants with previous experience with psychedelics completed all study visits, but only 17 (11 males, mean age = 33.8 y, SD = 8.3) had reliable ECG data (i.e., less than 10% noisy segments). Further details about the experimental procedure and participants can be found in Timmermann et al. (2023) (31).

**Figure 2.**
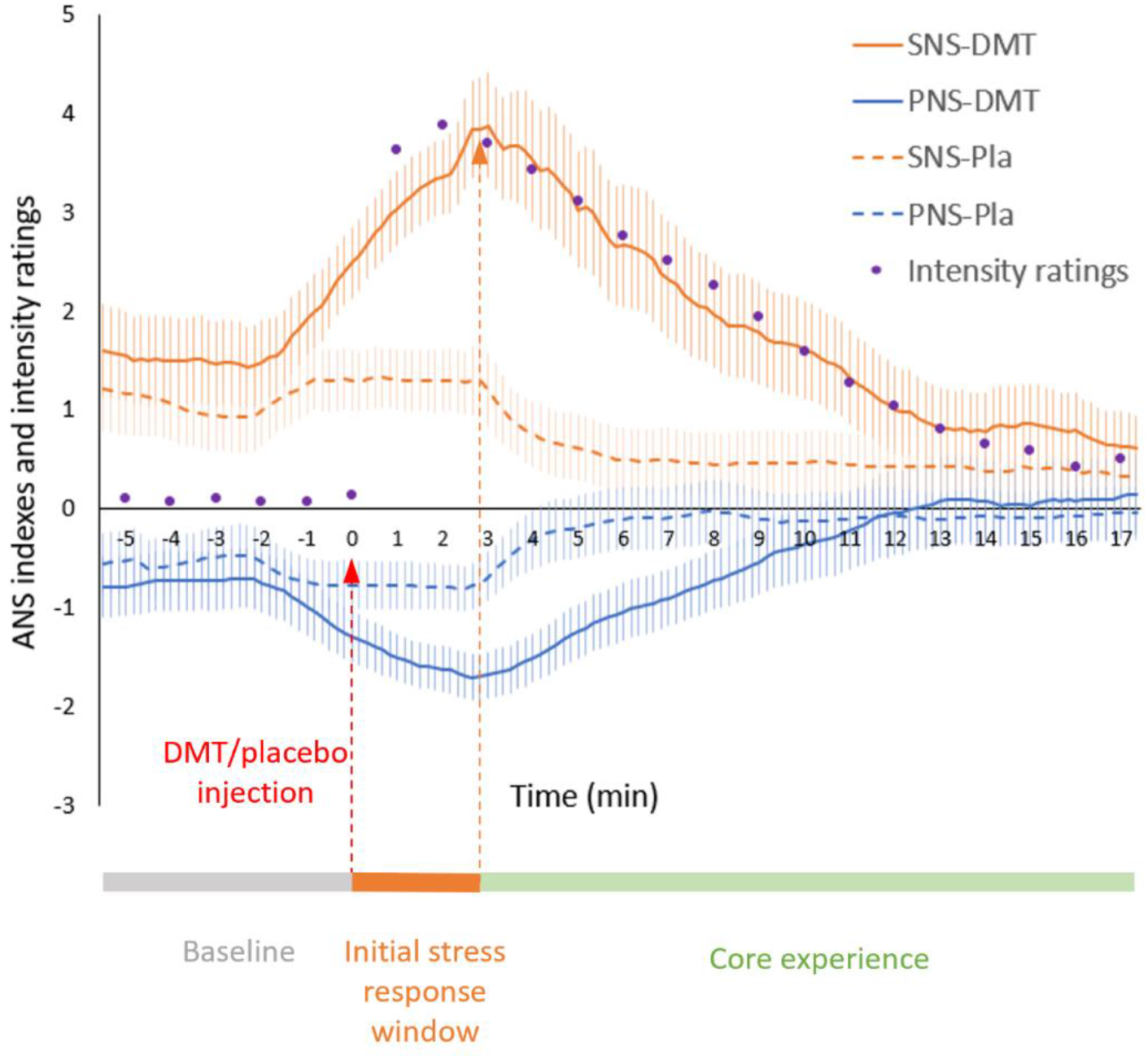
Measures of SNS (orange) and PNS (blue) activity, estimated based on 180 s of HRV data, with 20 s increments, were averaged across the 17 participants, for the DMT (continuous lines) and the placebo sessions (dotted line). Error bars indicate standard errors. Intensity ratings (purple dots), corrected within the same participants but during a distinct DMT session, were averaged across participants and normalized to match the levels of SNS indexes for visualization purpose. Significant differences between DMT and placebo can be found on Table S1. The line at the bottom illustrates the different phases which were considered for data handling. ECG data used to estimate SNS and PNS was collected during 28 min, with 8 minutes of baseline, followed by DMT/placebo injection. SNS and PNS indexes were computed from 180 s of data, therefore no measures could be computed for the first and last 3 minutes of ECG data. The onset zone, or initial stress response, was defined as the phase during which SNS ascents. Note that for both DMT and placebo, SNS starts increasing before the injection, most likely due to anticipatory processes.

This study was approved by the National Research Ethics Committee London—Brent and the Health Research Authority and was conducted under the guidelines of the revised Declaration of Helsinki (2000), the International Committee on Harmonization Good Clinical Practices guidelines, and the National Health Service Research Governance Framework. Imperial College London sponsored the research, which was conducted under a Home Office license for research with Schedule 1 drugs.

### Subjective ratings

Participants rated their subjective experience at the end of their DMT or placebo session, using the 11 Dimensions Altered States of Consciousness Questionnaire—ASC-11D (31). Among the 11 subscales of the ASC-11D, our analysis focused on those most relevant to evaluate the positive characteristics of peak experiences - *Experience of Unity, Spiritual Experience, Blissful State, Insightfulness* – and the elements of the experience that carry a negative valence - *Impaired Control and Cognition* and *Anxiety*. We also reported the results for the five other subscales (*Disembodiment, Complex Imagery, Elementary imagery, Synaesthesia* and *Meaning*) in order to control for the specificity of the effects observed, but did not take these into account in our multiple comparisons corrections. Participants also completed the Well-being index (WHO-5) questionnaire at baseline and two weeks after their experience (34).

### ECG recording

#### Data collection

ECG data was collected using two electrodes. One was placed in participants’ backs (behind the chest area), and the other was placed above the heart area. The data was recorded with an MR-compatible BrainAmp MR amplifier (BrainProducts GmbH, Munich, Germany). The data was recorded with a sampling rate was 5 kHz, and with a hardware 250 Hz low-pass filter. Recordings lasted 28 min, with 8 min of baseline and 20 min post injection.

#### Pre-processing

ECG data was demeaned and band-passed filtered at 1-30 Hz. It was then exported in text format to Kubios Scientific software (35). Automatic R-peak detection was applied, and all identified R-peaks were again manually inspected. Where necessary, R-peaks were corrected manually. Data sets with >10% of noisy segments (i.e. without the possibility of reliable R-peak detection due to artifacts) were discarded from further analysis. Three participants for the DMT session and four participants – same three plus another – for the placebo session were discarded due to this procedure.

### Measures of ANS activity

A similar procedure to that used in Olbrich et al. was used, whereby indexes of sympathetic (SNS) and parasympathetic (PNS) activity were computed in Kubios (18) based on 180s RR-intervals data, updated every 20s. To evaluate the relationship between ANS measures and subjective ratings during the core experience, we focused on the window following the initial stress response (or the ‘onset’), when SNS activity is beginning to reduce (3 minutes post injection, at minute 11, Figure 2), and during which subjective intensity ratings are significantly superior to that of placebo (up to minute 25, Table S1). For the correlation analyses relating to specific periods (i.e. baseline or core experiences), the indexes were computed based on the corresponding section of RR interval data (3 to 7 min for baseline, 11 to 25 min for core experience, see Figure 2).

#### Measure of sympathetic activity

A Sympathetic Nervous System (SNS) index was calculated in Kubios using mean HR (higher heart rate is linked to higher sympathetic cardiac activation), Baevsky’s stress index (a geometric measure of HRV reflecting cardiovascular system stress, and sympathetic cardiac activation) and standard deviation along the line of identity of the R-R Pointcaré plot (SD2, see below) (35).

#### Measure of parasympathetic activity

A Parasympathetic Nervous System (PNS) index was calculated in Kubios based on mean R-peak intervals (RR) (longer mean RR interval means lower heart rate and higher parasympathetic cardiac activation), the root mean square of the successive differences (RMSSD) - which is a commonly used time-domain HRV parameter that captures the quick beat-to-beat changes in RR interval, and therefore, strongly linked to respiratory sinus arrhythmia, a well-known measure of parasympathetic activity - and Poincaré plot index SD1 (35).

#### Measures of sympatho-vagal balance

Given the controversy around the validity of the ratio of low frequency (LF) to high frequency power (HF) - computed from Fast Fourier Transformation of R-R peaks time series - as an index of sympatho-vagal balance (36, 37), we used a measure derived from the PoinCarré plot of R-R intervals that has been proposed as an index of sympatho-vagal balance (38, 39). The Poincaré plot is a scatter plot of *RR*_*n*_ vs. *RR*_*n+1*_ where *RR*_*n*_ is the time between two successive R peaks and *RR*_*n+1*_ is the time between the next two successive R peaks. When the plot is adjusted by the ellipse-fitting technique, the analysis provides three indices: the standard deviation of instantaneous beat-to-beat interval variability (SD1), the continuous long-term R/R interval variability (SD2), and the SD1/SD2 ratio (SD12) (40). SD1 indicates the dispersion along the minor axis of the Poincaré plot’s fitted ellipse. It reflects short-term HRV (instantaneous beat-to-beat variability) and correlates with baroreflex sensitivity and is therefore mainly influenced by parasympathetic modulation. SD2 indicates the dispersion along the major axis of the Poincaré plot’s fitted ellipse and reflects long-term RR interval fluctuations. SD2 increases suggests activation of both parasympathetic and sympathetic nervous system. SD1/SD2, which measures the unpredictability of the RR time series, is used to measure autonomic balance (38).

#### Measure of sympathovagal coactivation

As we hypothesized that peak experiences would be associated with a state of co-activation of PNS and SNS, we computed a PNS and SNS interaction index based on the product of the two measures, previously translated into a range of strictly positive values. A higher SNSxPNS index indicates more co-activation of the two ANS branches.

### Statistical analysis

To visualize the time course of SNS and PNS activity during the DMT and placebo sessions, the indexes for each time-points were averaged across participants. Paired samples t-tests were used to identify the time points where SNS and PNS indexes were significantly different between the DMT and placebo sessions (Table S1).

After checking for the normality of each measure, Spearman correlations were used to assess the relationship between ANS activity and subjective experience, as measured using the ASC-11D questionnaire (Table 1). All significance levels were set to *p* < .05 (two-sided). Tables and figures show uncorrected statistics. Bonferroni corrections were only used for the correlations that were not predicted and are reported when relevant in the results section.

**Table 1.** Pearson correlations between SNS and PNS measures averaged during the core experience (post-ascent phase) and the subscales of the ASC-11D questionnaire. * p<0.05, and ** p<0.01, uncorrected

|  | **SNS** | **PNS** | **SNSxPNS** |
| --- | --- | --- | --- |
| **Unity** | $r=.517^*$ | $r=-.436$ | $r=.051$ |
| **Spiritual Exp** | $r=.07$ | $r=.155$ | $r=.683^{**}$ |
| **Blissful state** | $r=.065$ | $r=.086$ | $r=.386$ |
| **Insightfulness** | $r=.205$ | $r=.055$ | $r=.783^{**}$ |
| **Disembodiment** | $r=-.281$ | $r=.156$ | $r=-.188$ |
| **Impaired Cognition** | $r=-.563^*$ | $r=.263$ | $r=-.173$ |
| **Anxiety** | $r=-.497^*$ | $r=.334$ | $r=-.067$ |
| **Complex Imagery** | $r=-.168$ | $r=.150$ | $r=.281$ |
| **Elementary Imagery** | $r=.528^*$ | $r=-.467$ | $r=-.118$ |
| **Synaesthesia** | $r=-.279$ | $r=.100$ | $r=.098$ |
| **Meaning** | $r=-.269$ | $r=.129$ | $r=-.015$ |

Correlations between minute-by-minute ANS indexes and subjective ratings were also performed to identify the phases of the experience during which the relation between ANS and subjective experience were strongest. Spearman coefficients for each of the correlations were then plotted for ease of visualization. Only subscales that showed at least one significant correlation with one of the ANS measures were plotted (Figure 3).

**Figure 3:**
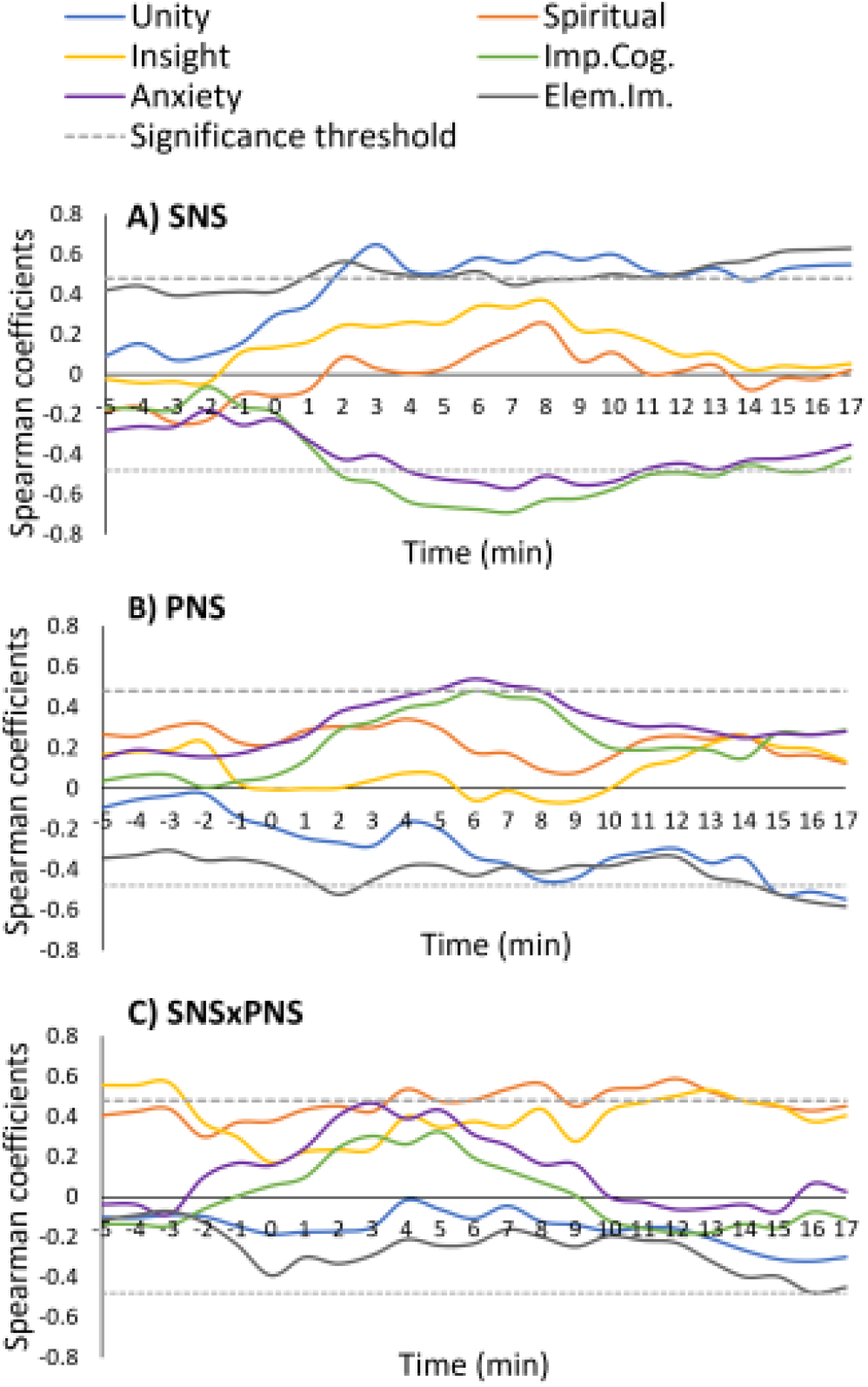
Evolution of correlations coefficients during the DMT experience. Spearman coefficients for the correlations between minute-by-minute measures of SNS (a), PNS (b) and SNSxPNS (c) are plotted over time. DMT was injected at minute 0. The cut off for correlation significance (i.e p<0.05) is indicated by the grey dotted lines and correspond to a r value of +/- 0.48. For clarity, only the scales showing some significant correlations with at least one of the 3 measures (SNS, PNS or SNSxPNS) were plotted.

Finally, participants were split into two groups based on the change in wellbeing reports from baseline to follow-up assessment at two weeks (WHO-5), using a median split. This procedure was used to account for participants inter-individual variabilities in estimating and reporting their wellbeing. Wellbeing data was missing for one participant. SNSxPNS scores during the core experience were compared between the two groups using an independent t-test.

## Results

### Effect of DMT vs Placebo on SNS and PNS activity profiles

The time courses of SNS and PNS activity, which were derived from ECG data collected during an 8 min baseline period and for 20 min post-DMT/placebo injection (see Methods) were averaged across participants during the DMT and the placebo session and plotted on **Figure 2**, along with during-experience subjective intensity ratings. SNS activity increased, and PNS activity decreased significantly in the DMT condition compared with placebo from the time of injection. SNS remained significantly higher in the DMT condition compared to placebo for 11 min post injection, and PNS remained significantly lower for 8 minutes (see **Table S1**).

### Association between ANS scores and subjective experience ratings

To the extent of our knowledge, there is no established index of sympatho-vagal coactivation. We therefore computed a new measure based on the product of the SNS and PNS indexes calculated in Kubios software based on ECG data during the core DMT experience (i.e. post-ascent or ‘core’ phase, after the initial physiological stress response) (see Methods). Importantly, and in keeping with our hypothesis, the SNS and PNS coactivation index, which we predicted would reflect both the quality and intensity of peak experiences, was positively and robustly correlated with *Spiritual Experience* (n=17, r=.625, p=0.007, uncorrected) and *Insightfulness* (n=17, r=.692, p=0.002, uncorrected) (Table 1), two important indicators of peak experiences.

SNS and PNS activity during the core experience showed some correlations with subjective ratings on the ASC-11D subscale. Spearman correlation coefficients are reported in **Table 1**. SNS activity during the core experience correlated positively with *Experience of Unity* (n=17, r=.517, p=0.034), and Elementary imagery (n=17, r=.528, p=0.035), and negatively with the two subscales of the ASC-11D of negative valence, *Impaired control and cognition* (r=-.590, p=0.013), and *Anxiety* (r=-.514, p=0.034), but these relations, which were not explicitly predicted, did not survive correction for multiple comparisons.

To get a more detailed view of which phases of the experience showed the strongest correlations between subjective ratings and ANS measures, Spearman correlations were also performed between the 11 subscales of the ASC-11D and minute-by-minute fluctuations of SNS and PNS indexes, calculated based on 180s RR-intervals data (**Figure 3**, Tables S2-S4). SNS was positively correlated with the *Experience of Unity* ratings from minute 2 to minute 17 post-injection and with *Elementary Imagery* from minute 1 to minute 17 post-injection. SNS was negatively correlated with *Impaired Cognition* from minute 2 to 15 post-injection and with *Anxiety* from minute 4 to 10 post-injection (**Figure 3a**, Table S2). PNS showed an inverted and weaker trend compared to SNS, with negative correlations with *Experience of Unity* from minute 15 to 17 and with *Elementary Imagery*, and positive correlations with *Anxiety* from minute 5 to 8 post-injection (**Figure 3b**, Table S3). Importantly, SNS x PNS was significantly and positively correlated with *Spiritual Experience from* minute 4 to 13 post-injection, and with *Insightfulness*, with correlations starting at baseline (minutes -5 to -3, maximum at minute -3), and resurfacing later in the experience at minutes 12 and 13 (maximum at minute 13) (**Figure 3c**, Table S4).

### Effect of autonomic balance on the quality of the peak experience

We hypothesized that peak experiences would be facilitated when the two branches of the ANS are well balanced at baseline. We therefore investigated the impact of autonomic balance at baseline on subjective ratings relevant to the peak experience and anxiogenic experience, using an established index of sympatho-vagal balance derived from the Poincaré plot of the R-R intervals, SD1/SD2 (38). SD1/SD2 at baseline significantly related to *Spiritual Experience* ratings (n=17, r=.555, p=0.021), and related marginally to *Insightfulness* (n=17, r=.413, p=0.099), but not to any other ratings (Table S5). Individuals who entered the DMT experience with a SD1/SD2 score closer to 1 (i.e., with a more balanced ANS), scored higher on the *Spiritual Experience* subscale of the ASC-11D (Figure S1). SNS and PNS indexes at baseline did not correlate with any elements of the subjective experience.

### Changes in ANS during the experience are related to long-term changes in wellbeing

Participants were split in two groups based on whether they had reported an improvement in their wellbeing in the two weeks following their DMT experience. Group 1 reported no change or a reduction in wellbeing (n=8, mean change in WHO-5 from baseline: -1.50 ± 1.6 SD). Group 2 reported improved wellbeing compared with baseline (n=8, mean change in WHO-5 from baseline: 1.62 ± 0.91). In terms of quality of the DMT experience, Group 2 showed significantly higher ratings of *Blissful Experience* compared to Group 1 (t=2.7, df=14, p=0.017), with no other significant differences for the other subscales. Importantly, when comparing ANS activity during the core experience, Group 2 showed a higher SNS x PNS coactivation index during the core experience compared with Group 1 (t=2.32, df=14, p=0.036), but no significant difference in SNS or PNS activity alone.

## Discussion

The aim of this study was to explore whether, and to what extent, changes in autonomic function during DMT administration related to the content of subjective experiences - as well as later changes in wellbeing. Our hypotheses were that positively experienced peak states would be associated with the coactivation of sympathetic and parasympathetic systems, while negatively valenced experiences would be associated with parasympathetic withdrawal. We also hypothesized that entering the experience in a state of greater sympatho-vagal balance would be conducive of peak experiences. While our results confirmed the involvement of both branches of the ANS in specific aspects of the peak experience, notably the *Spiritual Experience* and *Insightfulness*, negative experiences did not appear to be associated with PNS withdrawal. Rather, we unexpectedly observed that negative experiences were associated with reduced engagement of the SNS during the core experience.

Only one study to-date directly investigated the effect of a psychedelic (LSD) on cardiac sympathetic-vagal activity, based on measures of heart rate variability (HRV) (18). However, in this study, HRV was only measured during two relatively small time-windows, out of many hours of LSD-induced psychedelic experience, making it difficult to interpret their relevance to the overall subjective experience. Furthermore, the absence of baseline HRV measurement prevented the assessment of the relevance of the ANS in predicting the quality of the subsequent psychedelic experience. Another study examined the effects of several psychedelics on HRV(41), however, this study did not directly examine measures of SNS and PNS and its relationship to subjective effects.

The present findings confirmed our hypothesis of the involvement of both PNS and SNS in peak experiences. More specifically, we found these effects to be more pronounced in the *Spiritual Experience* and *Insightfulness* subscales, both features of peak states induced by psychedelics (31). Importantly, sympatho-vagal coactivation during the experience was a better predictor of subsequent improvement in wellbeing than subjective reports. Future work should evaluate whether the present findings are specific to peak experiences induced by DMT or if they can be extended to other psychedelics and non-drug induced NSCs. Such a physiological marker of peak experience would allow the implementation of techniques aimed at guiding individuals undergoing psychedelic-assisted therapy towards favorable physiological states via biofeedback, where behavior (e.g., breathing) is guided by real-time feedback of the ANS. Furthermore, these physiological peak states could serve as objective markers of the quality of the psychedelic experience, which could be particularly valuable given the ineffability inherent to peak experiences.

It is important to note that, although the present results confirm our hypothesis that sympatho-vagal coactivation is an optimal physiological state for the occurrence of psychedelic-induced peak experience, SNS and PNS can also coactivate under other circumstances that are not related to such experiences, such as the ‘freezing’ state (42), or certain forms of panic disorder, whereby increased excitability of both the PNS and SNS, referred to as amphotonia, can occur (43). Possibly accounting for this apparent discrepancy, according to a popular theory, the PNS may be further divided into two sub-systems - a ventral one associated with rest, safety and pro-social behaviors, and a dorsal one associated with immobilization behaviors (44).

The observation that baseline sympatho-vagal balance can predict the intensity of subsequent spiritual experiences is also of interest, as it may provide a physiological marker of ‘readiness’ to undergo a deeper and more transformative psychedelic experience. It is also consistent with the notion, pursued in some contemplative practices such as Zen Buddhism, that finding the right equilibrium between arousal and rest, tension and relaxation, is key to the meditative practice (45). If indeed meditation practices allow the cultivation of a more balanced ANS, as suggested in a study that identified increase sympatho-vagal balance after an advanced meditation program (46), our findings may provide mechanistic evidence for previous observations that contemplative practices may aid in the preparation for psychedelic experiences to foster safety and positive effects (1, 47). Our findings may thus have translational relevance by suggesting that cardio-physiological state may be a key factor for the success of the clinical use of psychedelics, complementing preliminary findings that psychological preparedness may promote safety and efficacy of psychedelic therapy (48). Furthermore, our findings are in line with suggestions that a cultivation of optimal psychological and physiological states can improve safety and efficacy of psychedelics and non-ordinary states of consciousness, more broadly (1, 49). These results warrant further research into the potential to develop markers of ‘readiness for acute alterations of consciousness’ based on the state of the ANS, which could be used as predictors of peak experiences, and thereby of positive outcomes in psychedelic-assisted therapy (4, 5).

DMT induced a pronounced increase in SNS activity, with a profile that appeared to directly parallel that of subjective intensity ratings (Figure 2). Though this association could simply relate to greater target receptor engagement by DMT other explanations are possible. Indeed, other forms of non-ordinary states of consciousness (NSCs) are also known to be associated with, and even induced by, states of high arousal. For instance, direct manipulation of the SNS through specific breathing practices (i.e., hyperventilation) can induce NSCs similar to those induced by psychedelics (50, 51). Noteworthily, near-death experiences, which share commonalities with DMT experiences (52), are associated with intense sympathetic activation, due to hypoxia (53).

Unlike the non-specific relations between SNS activity and numerous aspects of the psychedelic experience previously reported with LSD (18), in this study, we identified a more specific relationship between SNS activity and psychedelic phenomenology, with a positive relationship principally observed for the ASC-11D subscale *Experience of Unity*. Like the *Spiritual Experience* and *Insightfulness* subscales, the *Experience of Unity* subscale is a feature of psychedelic peak experiences (31). It comprises several items indicating a disruption to the sense-of-self, including feelings of time and space collapsing and merging with the environment. Notably, states of high arousal induced via other means have also been found to disrupt time and space processing (54) – possibly as a result of a disconnection of the hippocampus with prefrontal areas under stress (55) - which may impact the ability to construct spatially coherent internal scenes and a sense of separate self (56). Relatedly, the role of endogenous serotonin transmission in non-drug induced NSCs has also been considered and discussed (57).

Although we were able to identify clear relationships between acute changes in ANS and subjective experiences elicited by DMT, the question of whether changes in the ANS are driving or following changes at the level of the central nervous system remains open. This could be addressed in future studies by comparing the time courses of ANS fluctuations with that of brain activity determined via electroencephalography or functional magnetic resonance imaging, in an approach similar to that of Candia-Rivera et al. (13). While we did not address ANS-CNS causality, we did find a prediction of positive peak experiences via baseline ANS activity, which suggests a causal link from the ANS to phenomenology. Future studies should also aim to complement measures of autonomic activity derived from cardiac activity with other autonomic measures such as electrodermal activity, pupillometry, or piloerection.

The present results support the hypothesis that the remarkable psychological effects of psychedelics are not only the result of cortical neuronal mechanisms but involve multiple bodily systems - including the ANS. Specifically, we found that physiological stress response, as well as the balance between physiological stress and relaxation, are key aspects of both peak experiences and improved wellbeing. These findings may help the development of biofeedback techniques to guide the ANS toward states that are conducive to experiences that facilitate better mental health outcomes.

## Acknowledgments

This study was funded via a donation by Patrick Vernon (mediated by The Beckley Foundation). V.B. is funded by the Beckley Foundation. C.T. is funded by Comisión Nacional de Investigación Científica y Tecnológica and Anton Bilton. R.C.-H. was funded by the founding funders of the Centre for Psychedelic Research and is now supported by a Ralph Metzner endowment.

