## Supplemental Information for "Autonomic Nervous System activity correlates with peak experiences induced by DMT and predicts increases in wellbeing"

**SUPPLEMENTAL INFORMATION (SI)**

SI TABLES

### Table S1

P values for the paired sample T-test comparing SNS, PNS and intensity measures between DMT and placebo minute by minute

| Time relative to injection (min) | SNS | PNS | Intensity ratings |
|---|---|---|---|
| -5 | 0.818 | 0.764 | 0.083 |
| -4 | 0.587 | 0.852 | 0.301 |
| -3 | 0.417 | 0.726 | 0.332 |
| -2 | 0.495 | 0.584 | 0.163 |
| -1 | 0.115 | 0.205 | 0.332 |
| 0 | 0.001 | 0.032 | 0.104 |
| 1 | <0.001 | 0.005 | <0.001 |
| 2 | <0.001 | 0.003 | <0.001 |
| 3 | <0.001 | 0.002 | <0.001 |
| 4 | <0.001 | 0.004 | <0.001 |
| 5 | <0.001 | 0.006 | <0.001 |
| 6 | <0.001 | 0.008 | <0.001 |
| 7 | 0.002 | 0.020 | <0.001 |
| 8 | 0.002 | 0.039 | <0.001 |
| 9 | 0.003 | 0.355 | <0.001 |
| 10 | 0.006 | 0.634 | <0.001 |
| 11 | 0.029 | 0.926 | <0.001 |
| 12 | 0.127 | 0.737 | 0.003 |
| 13 | 0.178 | 0.540 | 0.002 |
| 14 | 0.099 | 0.620 | 0.004 |
| 15 | 0.096 | 0.544 | 0.003 |
| 16 | 0.126 | 0.546 | 0.007 |
| 17 | 0.195 | 0.527 | 0.026 |

**Table S2**

Spearman correlation coefficients for the correlations between SNS activity minute by minute and subjective experience scores (11D-ASC). * and **indicate $p<0.05$ and $p<0.01$ respectively. Only r values for which $p<0.1$ are shown.

| Time relative to injection (min) | Unity | Spiritual | Blissful state | Insight. | Disembod. | Impaired Cognition | Anxiety | Complex Imagery | Elementary Imagery | Synaetshesia | Meaning |
|---|---|---|---|---|---|---|---|---|---|---|---|
| -5 | | | | | | | | | .420 | | |
| -4 | | | | | | | | | .442 | | |
| -3 | | | | | | | | | | | |
| -2 | | | | | | | | | | | |
| -1 | | | | | | | | | .413 | | |
| 0 | | | | | | | | | .413 | | |
| 1 | | | | | | | | | .491* | | |
| 2 | .525* | | | | | -.513* | -.426 | | .563* | | |
| 3 | .647* | | | | | -.546* | | | .518* | | |
| 4 | .515* | | | | -.423 | -.640** | -.490* | | .494* | | |
| 5 | .510* | | | | | -.663** | -.526* | | .486* | | |
| 6 | .581* | | | | | -.676** | -.542* | | .514* | | |
| 7 | .556* | | | | | -.690** | -.573 | | .445 | | |
| 8 | .608* | | | | | -.628** | -.508* | | .472 | | |
| 9 | .571* | | | | | -.620** | -.554* | | .472 | | |
| 10 | .598* | | | | | -.574* | -.537* | | .499* | | |
| 11 | .517* | | | | | -.504* | -.473 | | .484* | | |
| 12 | .493* | | | | | -.492* | -.446 | | .504* | | |
| 13 | .529* | | | | | -.508* | -.477 | | .550* | | |
| 14 | .468* | | | | | -.453 | -.429 | | .568* | | |
| 15 | .525* | | | | | -.486* | -.422 | | .614** | | |
| 16 | .542* | | | | | -.481 | | | .622** | | |
| 17 | .547* | | | | | -.416 | | | .629** | | |

**Table S3**

Correlations PNS activity minute by minute and subjective experience scores (11D-ASC). * indicates p<0.05 and p<0.01 respectively. Only r values for which p<0.1 are shown.

| Time relative to injection (min) | Unity | Spiritual Experience | Blissful state | Insight. | Disembod. | Impaired Cognition | Anxiety | Complex Imagery | Elementary Imagery | Synaesthesia | Meaning |
|---|---|---|---|---|---|---|---|---|---|---|---|
| -5 | | | | | | | | | | | |
| -4 | | | | | | | | | | | |
| -3 | | | | | | | | | | | |
| -2 | | | | | | | | | | | |
| -1 | | | | | | | | | | | |
| 0 | | | | | | | | | | | |
| 1 | | | | | | | | | -.440 | | |
| 2 | | | | | | | | | -.523* | | |
| 3 | | | | | | | .417 | | -.452 | | |
| 4 | | | | | | | .457 | | | | |
| 5 | | | | | | .422 | .492* | | | | |
| 6 | | | | | | .480 | .540* | | -.430 | | |
| 7 | | | | | | .450 | .508* | | | | |
| 8 | -.456 | | | | | .429 | .481 | | | | |
| 9 | | | | | | | | | | | |
| 10 | | | | | | | | | | | |
| 11 | | | | | | | | | | | |
| 12 | | | | | | | | | | | |
| 13 | | | | | | | | | -.435 | | |
| 14 | | | | | | | | | -.462 | | |
| 15 | -.520* | | | | | | | | -.521* | | |
| 16 | -.510* | | | | | | | | -.560* | | |
| 17 | -.547* | | | | | | | | -.582* | | |

**Table S4**

Correlations SNSxPNS minute by minute and subjective experience scores (11D-ASC). *indicates $p<0.05$ and $p<0.01$ respectively. Only r values for which $p<0.1$ are shown.

| Time relative to injection (min) | Unity | Spiritual Experience | Blissful state | Insight. | Disembod. | Impaired Cognition | Anxiety | Complex Imagery | Elementary Imagery | Synaesthesia | Meaning |
|---|---|---|---|---|---|---|---|---|---|---|---|
| -5 | | | | .556* | | | | | | | |
| -4 | | .424 | | .556* | | | | | | | |
| -3 | | .437 | | .568* | | | | | | | |
| -2 | | | | | | | | | | | |
| -1 | | | | | | | | | | | |
| 0 | | | | | | | | | | | |
| 1 | | .433 | | | | | | | | | |
| 2 | | .451 | | | | | | | | | |
| 3 | | .424 | | | | | | | | | |
| 4 | | .535* | | | | | | | | | |
| 5 | | .475 | | | | | | | | | |
| 6 | | .482* | | | | | | | | | |
| 7 | | .536* | | | | | | | | | |
| 8 | | .563* | | .437 | | | | | | | |
| 9 | | .450 | | | | | | | | | |
| 10 | | .531* | | .429 | | | | | | | |
| 11 | | .543* | | .470 | | | | | | | |
| 12 | | .587* | | .503* | | | | | | | |
| 13 | | .522* | | .531* | | | | | | | |
| 14 | | .477 | | .475 | | | | | | | |
| 15 | | .450 | | .455 | | | | | | | |
| 16 | | .428 | | | | | | | -.477 | | |
| 17 | | .451 | | | | | | | -.450 | | |

**Table S5:** Correlation between measures of sympathovagal (i.e., SNS-PNS) balance at baseline and during core experience and subjective ratings of the quality of the peak experience. * indicates p<0.05

| | **LF/HF Baseline** | **SD1/SD2 Baseline** |
|---|---|---|
| **Experience of Unity** | r=.176 | r=-.097 |
| **Spiritual Experience** | r=-.172 | r=.555* |
| **Blissful state** | r=-250 | r=.126 |
| **Insightfulness** | r=-.249 | r=.413 |
| **Impaired Cog.** | r=-.047 | r=-.177 |
| **Anxiety** | r=.248 | r=-.190 |

SI FIGURE

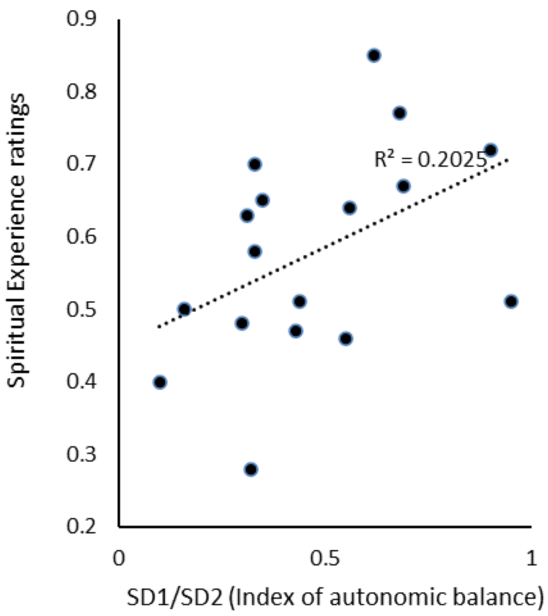


**Figure S1: Correlation between autonomic balance at baseline and Spiritual Experience ratings**

The x axis corresponds to the ratio of SD1 to SD2, measures derived from the Poincaré plot of the RR-intervals (see SI Methods). The y axis corresponds to individual scores of the ASC-11D subscale ‘Spiritual Experience’.
